# *In vitro* model reveals structural and metabolic insights to the porcine caecal microbiota in response to β-mannan exposure

**DOI:** 10.64898/2026.01.22.701113

**Authors:** Jenny Merkesvik, Lars J. Lindstad, Özgün C. O. Umu, Ronja M. Sandholm, Sabina L. La Rosa, Torgeir R. Hvidsten, Phillip B. Pope, Bjørge Westereng

**Author notes:** **Reason for author order:** based on contribution. JM and LJL contributed equally to the research and wrote the original draft. JM led the preparation of the manuscript, supplementary files, and raw data for submission. ÖCOU coordinated the in vivo animal trial. RMS contributed to bioinformatic data generation. SLLR, TRH, and PHP supervised the work. BW supervised the work and contributed to writing the original draft. TRH, PHP, and BW acquired funding for the research.

## Abstract

The gastrointestinal microbiota plays a pivotal role in shaping host physiology and health. By selectively promoting bacteria associated with improved host health, microbiota-directed fibres offer a strategy to enhance the beneficial functions of the microbiota. In this work, we developed a pH-controlled *in vitro* fermentation system (InVitSim) as a model to evaluate the effects of such a fibre – acetylated galactoglucomannan from Norway spruce – on the composition and functionality of porcine caecal microbial communities. We validated the experimental outcomes by comparing the response of the *in vitro* model to a previous *in vivo* feeding trial utilising the same β-mannan fibres. Long-read sequencing with Oxford Nanopore, metatranscriptomics, and short-chain fatty acid measurements were undertaken to survey microbial community dynamics and functionality. Microbial communities in pigs and InVitSim responded similarly to β-mannan supplementation, with taxa like *Prevotella, Catenibacterium*, and *Faecalibacterium* increasing in abundance. Intriguingly, some taxa were observed to be more affected by β-mannan supplementation in InVitSim than *in vivo*. These taxa included several bacterial species that were not previously known to utilise β-mannan, yet exhibited upregulated genes encoding carbohydrate-active enzymes involved in the degradation of this substrate.

**Importance:** In this study, we establish a fermenter system able to preserve more than 70% of over 300 distinct microbial taxa identified in the porcine caecal gut. The *in vitro* model and the functional omic data generated from it enabled us to identify relevant microbial populations that responded to the presence of AcGGM by upregulating β-mannan-specific polysaccharide utilisation loci. Our results highlight the value of *in vitro* approaches as a complementary tool to *in vivo* trials for learning about the gastrointestinal microbiome’s response to dietary interventions on the host level.

**Description of supplementary files:**

A. In-depth analyses for *in vitro* model validation and investigations.
B. Common taxa between *in vivo* and *in vitro* systems exposed to β-mannans.
C. Differential abundance analysis result visualisations.
D. Short-chain fatty acid concentrations and correlation with microbial abundances.

## Introduction

The gastrointestinal microbiota is essential for the health and well-being of mammals. In pigs, it helps maintain and regulate the immune system; defends against pathogens; and breaks down indigestible feed components like fibres that can be converted into host-beneficial short-chain fatty acids (SCFAs) (1,2). In swine production, health issues like infections, diarrhoea, and dysentery have been linked to gut microbiota dysbiosis (3–5). Disease susceptibility is particularly high when the gut microbiota is drastically changing during the weaning and post-weaning phases (6). These illnesses have detrimental effects on both the animals and the production process. Moreover, the use of antibiotics as growth promoters in animal feed has been banned in Europe effective from 2006 (Regulation (EC) No. 1831/2003), and the use of antibiotics in general is discouraged to mitigate the risk of antibiotic resistance (7,8). However, reduced antibiotic use may lead to higher disease frequency and mortality rate (9). Finding alternative methods to improve animal welfare and production efficiency has therefore become essential to meet our increasing resource demands. A central focus of this effort is to expand our understanding of the swine gut microbiome, its implications for host health, and how this knowledge can be applied to promote beneficial microbes.

Microbiota-directed fibres (MDFs) and prebiotics can be administered to production animals to improve gut health (10,11). These MDFs include carbohydrates that are selectively utilised by health-promoting microbes. β-Mannans are potential MDFs that have been shown to promote the growth of bacteria linked to positive health effects. Examples include important butyrate producers like *Roseburia* and *Faecalibacterium spp*., and several commensal Bacteroidota species (12–17). β-Mannan fibres are composed of a backbone of β-1,4-D-mannopyranose residues, which can be intersected with β-1,4-D-glucopyranose residues to form glucomannan. Additional decorations with α-1,6-D-galactopyranose residues yield galactomannan and galactoglucomannan (18). Furthermore, these β-mannans can be acetylated to various extents at the 2-*O*-, 3-*O*-, and 6-*O*-positions on mannose moieties. The fibres’ structural complexity necessitates multiple carbohydrate-active enzymes (CAZymes) for their breakdown. Complete degradation of complex β-mannans, such as acetylated galactoglucomannan (AcGGM), requires several glycoside hydrolases (GHs) and acetyl carbohydrate esterases (CEs) (19). The enzymatic machinery needed to metabolise β-mannans may be exploited to target and selectively promote beneficial bacteria that possess these enzymes.

*In vitro* experiments have been widely used to test the effects of different fibres on both single microbial species and communities of varying complexity. *In vitro* studies mitigate ethical, financial, and infrastructural challenges often associated with *in vivo* experiments. They can provide insight into substrate degradation and utilisation, microbial interactions, and metabolite production (e.g., SCFAs) (20,21). Several single- and more complex multi-compartment *in vitro* models have been developed in recent years to simulate and study the gastrointestinal ecosystem of mammals (17,20,22,23). Unfortunately, the versatility of such systems is often diminished due to a limited representativeness of *in vitro* microbial communities compared to their *in vivo* counterparts. For instance, a recent paper presented a 15% concordance between their multi-compartment model and donor pigs (22). A likely explanation for the loss of species richness reported in several *in vitro* studies is the long-term adaptation and stabilisation of the microbiota to *in vitro* conditions (22,24).

We conducted an *in vitro* fermentation experiment inoculated with caecal material from pigs after a 4-week *in vivo* feeding trial with Norway spruce (*Picea abies*) AcGGM supplements (**Fig. 1**). This study was designed to minimise the loss of species richness during *in vitro* cultivation by (a) focusing on processing faecal material in ways that preserved the highest possible viability of bacterial taxa, and (b) sampling at time points when the majority of bacteria were in exponential growth during fermentation. Through this approach – which we refer to as InVitSim – we retained over 70% of 325 taxa present in the *in vivo* samples. We validated the effect of AcGGM on the microbial communities in the *in vitro* fermentation setup by using metagenomic data to track changes in the *in vitro* population abundances, which we compared with data from another *in vivo* feeding trial. Secondly, we used functional data in the form of metatranscriptomics to investigate gene expression patterns of the enzymatic machinery required to degrade AcGGM fibres. Measured SCFA levels further tied the observed community developments to AcGGM degradation by members of the porcine caecal microbiota. We propose new potential AcGGM degraders based on their increased abundance and expression levels of polysaccharide utilisation loci (PULs) when AcGGM was supplemented *in vitro*.

**Figure 1.**
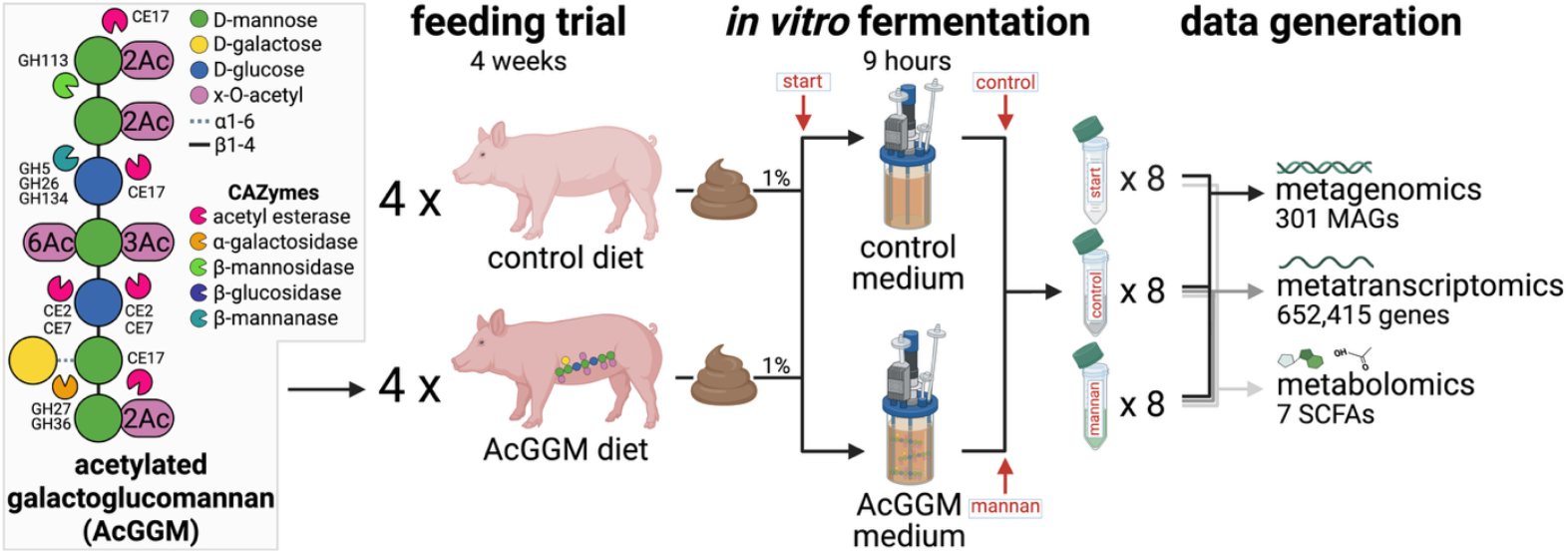
The experimental setup spanned a 4-week animal trial with acetylated galactoglucomannan fibre supplementation and a subsequent in vitro fermentation using growth medium with or without added fibre. The 24 collected samples thus comprised 8 pre-fermentation (“start”) samples from either porcine diet group, and 16 post-fermentation samples – 8 from each porcine diet group – grown in medium with (“mannan”) or without (“control”) fibre supplements. Three omic layers were generated from these samples: metagenomics, metatranscriptomics, and targeted metabolomics covering select short-chained fatty acids. Figure created in BioRender; Merkesvik J (2025), https://biorender.com/o06u268.

## Results and Discussion

In this study, we investigated the porcine gut microbiome and its functional capability for degrading complex carbohydrates in response to AcGGM administration using InVitSim; a fermentation approach for modelling the *in vivo* porcine caecal microbiome. Following an *in vivo* feeding trial where pigs were fed either an AcGGM-supplemented or an AcGGM-free (referred to as *control*) diet, caecal digesta samples were inoculated in fermentors with medium with or without added AcGGM as a carbon source. A total of 24 samples across six different sample groups were collected from InVitSim. For each of the porcine diets – control (first letter **c**) and AcGGM (first letter **m**) – there were three samples: before fermentation (**cs, ms**); after fermentation in medium without added carbon sources (referred to as control growth medium) (**cc, mc**); and after fermentation in an AcGGM-supplemented medium (**cm, mm**) (**Fig. 1**).

We first established the performance of the fermentors by how well our *in vitro* model replicated population dynamics observed in the *in vivo* porcine gut microbiome. InVitSim was therefore benchmarked against observations from Michalak *et al*. (13). Briefly, this paper from 2020 describes a 28-day pig feeding trial where one group was given AcGGM supplements at a 4% inclusion level; the same as in the present trial. Secondly, we investigated the effect of AcGGM on the porcine caecal microbiome in the present trial. Specifically, we assessed the microbial community composition, functional capability, gene expression patterns, and SCFA levels associated with *in vivo* and *in vitro* fibre administration.

### Benchmarking: how well does InVitSim reflect the *in vivo* experiments?

To assess the reliability of the InVitSim approach as a model for the *in vivo* porcine caecal microbiome, we studied the microbial community structure in samples from animals given an AcGGM-free (control) diet during the four-week feeding trial. With these samples, we compared observations from before and after fermentation in growth medium with either AcGGM supplementation or control medium without any added carbohydrate source.

Considering all identified metagenome-assembled genomes (MAGs) with completeness over 50% and contamination under 5%, the inoculated samples from pigs given the control diet contained an average of 325 distinct populations. From samples taken from fermentors in the exponential phase (9 hours) in either control or AcGGM-supplemented media, we saw a 25% and 29% decrease in species richness, respectively, averaging 246 and 232 populations still present after *in vitro* fermentation (**Fig. 2A**). Shannon entropy diversity – a nonlinear index which considers population abundances (25) – reported a similar diversity decrease from the pre-to the post-fermentation stage. Accounting for the microbiota’s phylogenetic relationships and functional capabilities, we saw that the post-fermentation state appeared more diverse than the original samples; with microbial communities inoculated in a control medium having experienced the strongest increase.

**Figure 2.**
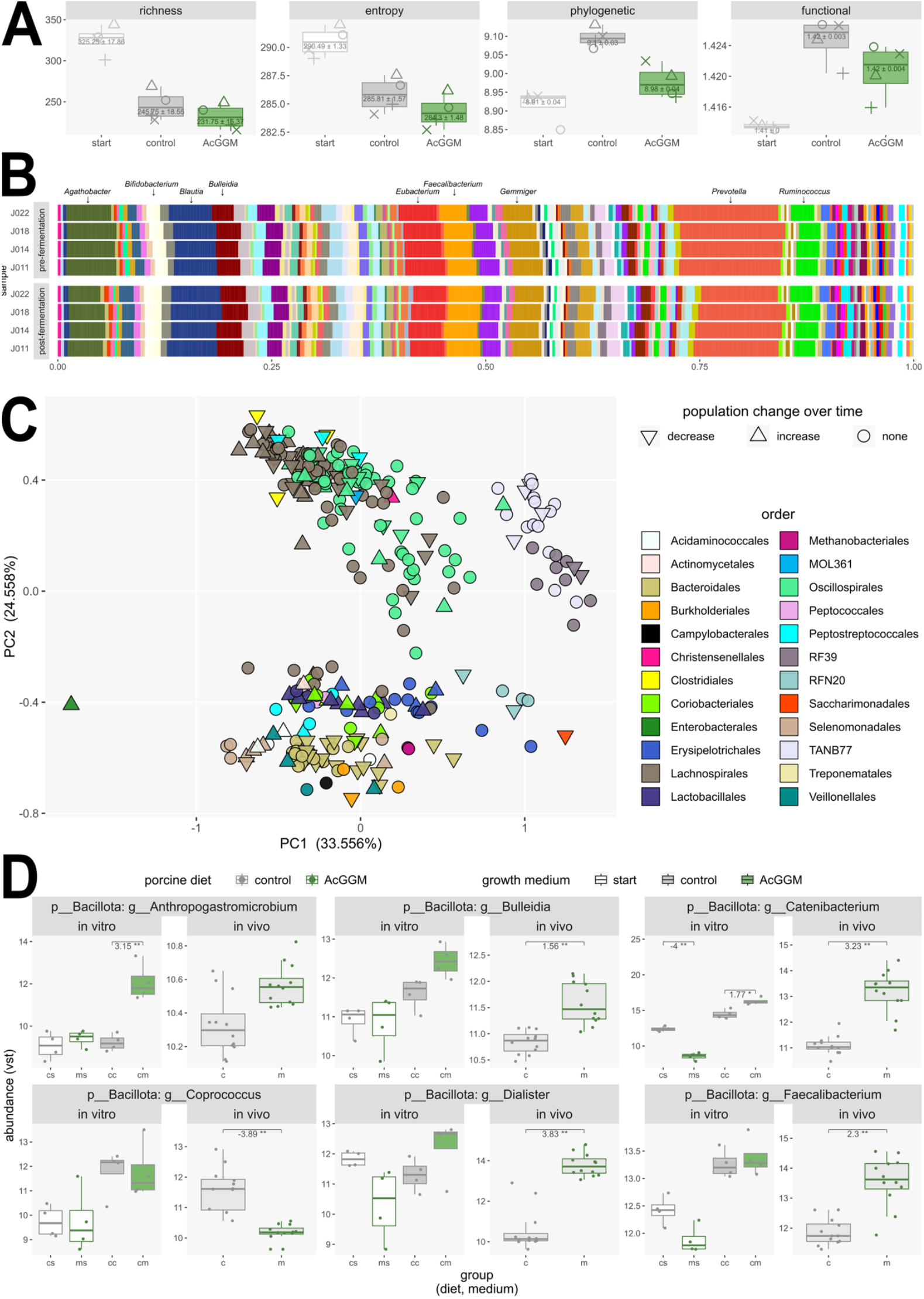
Investigations of the microbial communities following fermentation in InVitSim. **A)** Alpha diversity measures in samples from pigs fed control diets before fermentation (“start”) and after 9 hours in different growth media (“control”, “AcGGM”) in InVitSim. Numbers overlaying each box denote the mean and standard deviation across the samples in the respective group. Facets from left to right display; species <u>richness</u>, measured in counts of unique populations present; Shannon <u>entropy</u> diversity index, scaling species richness by observed abundances; <u>phylogenetic</u> diversity, based on a phylogenetic tree of the MAGs; and <u>functional</u> diversity, based on genome-inferred functional traits derived from KEGG ontology annotations. **B)** Relative abundances of genera in samples from control diet pigs, grouped according to whether the sample was extracted before or after the 9-hour in vitro fermentation. Each bar in the “post-fermentation” category comprises samples grown in control and AcGGM-supplemented media, leaving the sampling time point as the central contrast in this comparison. The full figure with legends is included in Suppl. A. **C)** Principal component analysis of the genome-inferred functional trait values for the 301 MAGs. Points cluster well according to taxonomic classification (colour) rather than according to whether the population was found in increased, decreased, or unchanged abundance over time (shape). The four most abundant genera in the in vitro fermentation (Agathobacter, Blautia, Eubacterium, and Prevotella) belong to the orders Lachnospirales (top left, brown cluster), Oscillospirales (top centre, mint green), and Bacteroidales (bottom centre, beige). **D)** Variance-stabilised abundances of select microbial populations at the genus-level of some common taxa between the present in vitro study and the 2020 in vivo study by Michalak et al. (16). Significant differential abundances between pairs of boxplots are indicated by horizontal bars with accompanying log2 fold change (thresholds absolute LFC>1, base mean>50) and FDR-adjusted p-values indicated by asterisks (*<0.05, **<0.01, ***<0.001). A full version with all common genera across in vitro and in vivo systems is included in Suppl. B. Abbreviations in axes: cs: control diet, before fermentation, ms: mannan diet, before fermentation, cc: control diet and medium, cm: control diet and mannan medium, mc: mannan diet and control medium, mm: mannan diet and medium.

To explain the patterns revealed by the alpha diversity metrics, we compared community compositions and their genome-inferred functional traits across samples before and after fermentation in InVitSim. These analyses are explained in detail in **Suppl. A**. Briefly, we found that the largest changes in abundance over time occurred within the already most numerous taxa (*Prevotella, Agathobacter, Eubacterium*) (**Fig. 2B**). Further, no taxa nor function was completely lost following fermentation (**Fig. 2C**). Taken together, our observations indicate that while the effective number of unique microbial populations in InVitSim decreased to around 70% of the original samples over time, the diminished populations overlapped strongly with the persisting microbial taxa in terms of both phylogeny and functional potential. Hence, InVitSim did not introduce any taxonomic or functional biases to the microbial community following *in vitro* fermentation.

Next, we aimed to determine whether the impact of AcGGM on the *in vitro* community reflected that of the *in vivo* caecal microbiota observed in the Michalak *et al*. study (13). Comparing alpha diversities for the two MAG catalogues (**Figs. 2A, SA2** in **Suppl. A**) revealed similar trends in reduced species richness and Shannon entropy from controls to AcGGM-supplemented samples. Thus, the fibre administration resulted in a narrower microbial community with fewer taxa (lower richness) of more even abundance (lower entropy) in both the *in vivo* and *in vitro* approaches. However, when looking at the phylogenetic diversity, the *in vivo* communities were not reduced when exposed to AcGGM as we observed in InVitSim. We ascribed phylogenetic diversity loss in our *in vitro* model to the reduced abundance of some of the community’s most dominating taxa – *Agathobacter, Eubacterium*, and *Prevotella* (**Fig. 2B, Suppl. A**) – however, this loss was not as evident *in vivo* (**Suppl. B**). Therefore, the observed difference in phylogenetic diversity between sample groups across the two study approaches may be attributed to InVitSim not catering as well to some prominent members of the gut microbiota as the *in vivo* setting, somewhat changing the community structure in disfavour of these functionally redundant populations. Despite this discrepancy, the InVitSim retained over 70% of 325 taxa in the *in vivo* samples, demonstrating its versatility compared to other *in vitro* methods in maintaining a relatively high species richness.

To find out whether the InVitSim microbiota responded similarly to AcGGM fibres as the *in vivo* microbiota, we compared population dynamics of taxa found in both the *in vitro* and *in vivo* MAG catalogues. We identified 104 overlapping species across 34 genera, and similar dynamics were observed in the two systems for populations with and without access to AcGGM for several genera (**Figs. 2D, SB2**). Specifically, the relative abundance patterns of these genera across the two diet groups pre-fermentation corresponded with the *in vivo* observations from Michalak *et al*. (13), and these patterns persisted after fermentation in media with or without AcGGM. Examples of these genera include *Anaerovibrio, Anthropogastromicrobium, Bulleidia, Coprococcus, Duodenibacillus, Enteromonas, Evtepia, Oribacterium*, and *Streptococcus*. We observed additional similarities between the InVitSim and *in vivo* communities’ response to AcGGM when only considering the relative population sizes in different growth media post-fermentation. Namely, data from both studies show genus-level population increases for *Paratractidigestivibacter, Catenibacterium, Dialister, Faecalibacterium, Holdemanella*, and *Mitsuokella* in AcGGM-supplemented samples; and population decreases for *Acetatifactor, Alloprevotella, Alitiscatomonas, Butyricicoccus, Dysmosmobacter, Faecousia, Floccifex, Lachnospira, Lactobacillus, Lawsonibacter, Limosilactobacillus*, and *Pararoseburia (****Suppl. B***).

Three of the genera with increased abundance following fibre supplementation in both InVitSim and the *in vivo* trial were highlighted in Michalak *et al*. as potential responders to AcGGM capable of producing SCFAs: *Catenibacterium* with acetate production; *Dialister* for propionate; and *Faecalibacterium* with production of both acetate and butyrate (13). *Prevotella* showed clear tendencies for genus-wide abundance increases during AcGGM supplementation only in InVitSim (**Fig. SB1**), but was also suggested as an AcGGM responder with propionate production (13). The remaining taxa named as potential SCFA-producing AcGGM responders in Michalak *et al*. – namely *Agathobacter* and *Megasphaera* – displayed no concise genus-wide AcGGM response in our *in vivo* nor *in vitro* approaches. While not regarded in the 2020 *in vivo* study, there were additional populations present in both InVitSim and our *in vivo* samples that appeared enriched when AcGGM was available. Examples include *Anthropogastromicrobium aceti* and several *Blautia* species (**Fig. SB2**). The potential AcGGM degradation activity of these taxa is explored in the next section.

As a final point on benchmarking the InVitSim approach against the *in vivo* porcine gut microbiota, we observed that some taxa appeared to thrive more in the *in vitro* setting by displaying a significant abundance increase post-fermentation. The genera *Anaerovibrio* (LFC>5), *Megasphaera* (LFC>4), *Oribacterium* (LFC>1), and *Escherichia* (LFC>7) (**Suppl. B**) all followed this trend. A notable decrease in population size was observed over time for *Agathobacter* (LFC<-4) and *Prevotella* (LFC<-2); however, no genera were completely depleted in InVitSim after fermentation. Likely explanations for these altered relative abundances in the *in vitro* setting are changes in interspecies competition, lack of the host’s influence both as a provider of habitats; nutrient exchange; and regulatory interactions across the host-microbiota barrier, and build-up of microbial metabolites. Additionally, possible effects related to the preparation of digesta samples for InVitSim inocula could contribute to *in vitro* and *in vivo* differences. Examples include low oxygen tolerance of some gut bacteria, prolonged storage of samples before fermentation, and the choice of growth medium (26–31). The basal growth medium used in this study is commonly employed in in vitro studies of the human and porcine gut microbiota.

In summary, the present *in vitro* approach for investigating the effect of AcGGM fibres on the porcine gut microbiota showed population developments for several key taxa (e.g., *Catenibacterium, Coprococcus, Dialister, Faecalibacterium*, and *Lactobacillus*) that were comparable to observations from an *in vivo* trial conducted more than four years earlier. Both *in vivo* and in our InVitSim approach, AcGGM-supplemented samples had lower species richness and entropy diversity than AcGGM-free control samples, indicating that the fibre supplementation led to a more specialised microbial community. Phylogenetic diversity was only different between control and AcGGM groups in the *in vitro* model, an observation attributed to the diminished presence of functionally and phylogenetically redundant populations from the community’s most dominating taxa (*Agathobacter, Eubacterium*, and *Prevotella*). While InVitSim did not appear to introduce any strong biases, some species seemed to experience an advantage when grown in the presence of AcGGM fibres *in vitro* (e.g., members of *Anthropogastromicrobium, Blautia*, and *Prevotella*). This observation underscores that, despite the concordance of 70% between InVitSim and *in vivo* samples and the overall well-preserved microbial community structure and function, there were some discrepancies when comparing population dynamics observed *in vitro* to those from an *in vivo* community exposed to the AcGGM fibres.

### Functional capabilities of the *in vitro* porcine gut microbiome

Having established that our InVitSim approach for modelling the porcine caecal microbiota yielded a similar development as seen in an *in vivo* fibre trial for several key taxa, we further investigated the microbiota’s response to AcGGM *in vitro*. In this section, we highlight populations, genes, and SCFAs that experienced significant changes in abundance, expression, and concentration, respectively, following AcGGM administration. In particular, we point out three AcGGM degraders from the InVitSim microbial community based on their increased population abundances and transcriptomic activity of proposed PULs when grown in the presence of AcGGM fibres. Additionally, we assessed the effect of prior *in vivo* exposure to the fibre by including microbial communities extracted from pigs fed an AcGGM-supplemented diet in the *in vitro* fermentation.

The present *in vitro* approach yielded a MAG catalogue of 301 taxa after filtering on completion (>50%), contamination (<5%), and presence in at least two samples to allow comparisons (**Fig. 3**). 299 MAGs were bacterial and spanned 23 orders and 122 genera, while two MAGs were classified in the archaeal order *Methanobacteriales*. The Bacillota orders Lachnospirales and Oscillospirales comprised over half of the unique populations in the catalogue (**Fig. 3C**). Each MAG contained an average of 2167 genes, and 400,855 of the total 652,415 genes were observed to be expressed in at least one of the collected samples. Considering the total number of transcriptomic reads mapped to each MAG, 53 populations experienced a significant expression level increase when AcGGM fibres were added to the growth medium in InVitSim (**Fig. 3A**). Examples include several *Eubacteria* and *Prevotella* that have previously been linked with SCFA production benefitting gut health (32,33).

**Figure 3.**
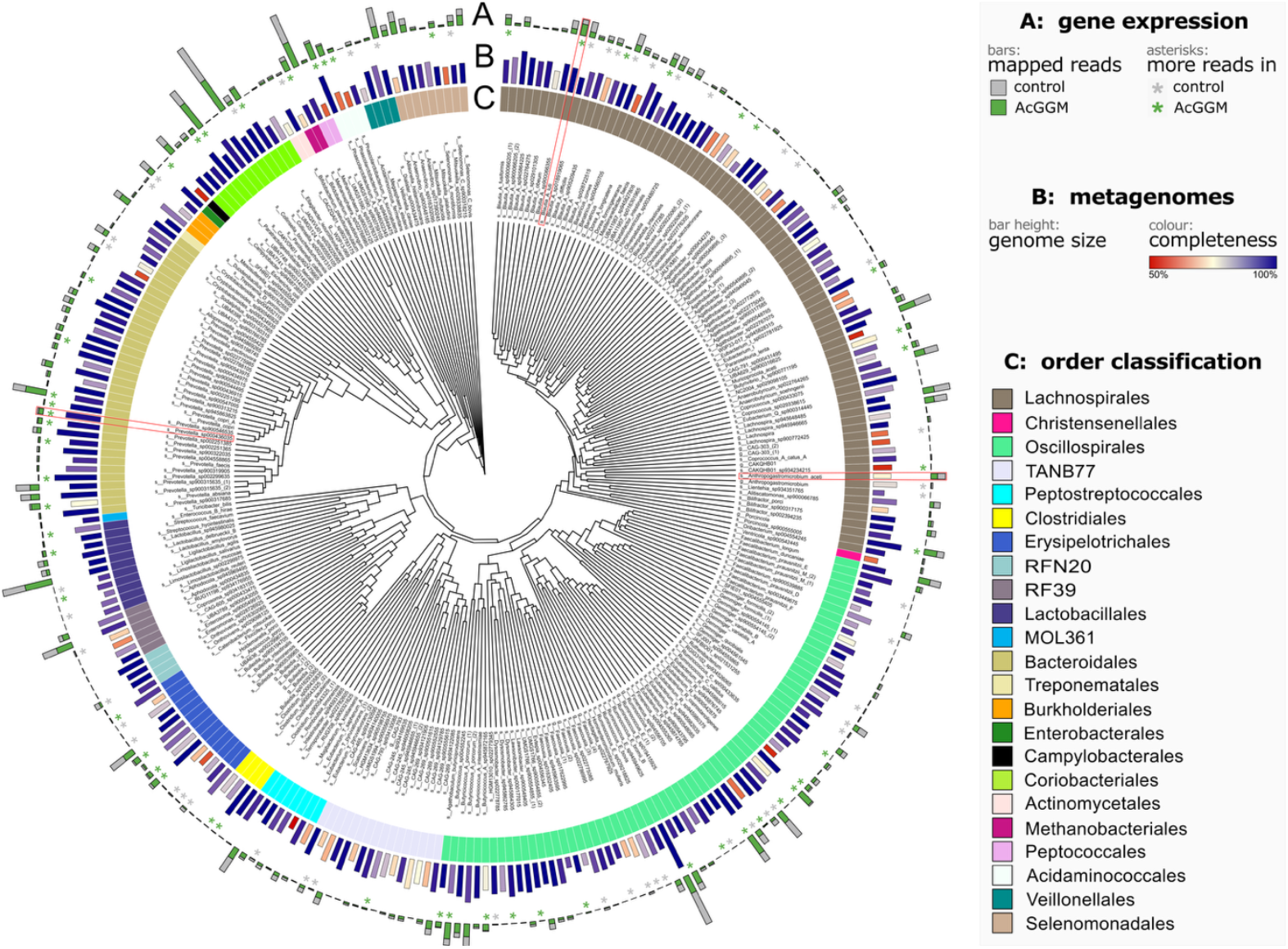
Phylogenetic tree of the 299 bacterial metagenome-assembled genomes of good quality identified in at least twice across the full sample set. The most specific taxonomic classifications are included as tip labels. Repeated taxon identifiers are numerated by order of appearance. Three populations of interest as AcGGM utilisers have been highlighted with red boxes: Anthropogastromicrobium aceti, Blautia luti, and Prevotella sp. **A)** Sum of variance-stabilised gene transcripts mapped to each genome across samples grown in either control or AcGGM-added medium. Asterisks mark populations with significant (absolute LFC>1, FDR p-value<0.05) differential expression between these groups, where asterisk colour indicates which growth medium yielded the most transcriptionally active population. **B)** Genome size represented by bar height, and MAG completeness relative to each respective reference genome indicated by colour. **C)** Taxonomic classification on the order level.

Among the microbial populations that appeared promoted in response to AcGGM supplementation in the InVitSim model were *Anthropogastromicrobium aceti, Blautia luti*, and *Prevotella sp000436035* (**Fig. 3**). These populations showed significant abundance increases when grown in the presence of AcGGM in InVitSim, but did not experience a similarly strong promotion following *in vivo* fibre supplementation (**Fig. 4**, left panels), where they seemed unable to compete with other better adapted microbes. Their increased abundance being related to fibre degradation was corroborated by elevated expression levels for several genes located in genomic regions representing PULs (**Fig. 4**, right panels). Moreover, upregulation of genes needed for mannan degradation occurred in AcGGM-grown populations from both controls and AcGGM-fed animals. These populations likely persisted by utilising alternative nutrient sources and only activated their mannan-degrading machinery upon exposure to AcGGM *in vitro*.

**Figure 4.**
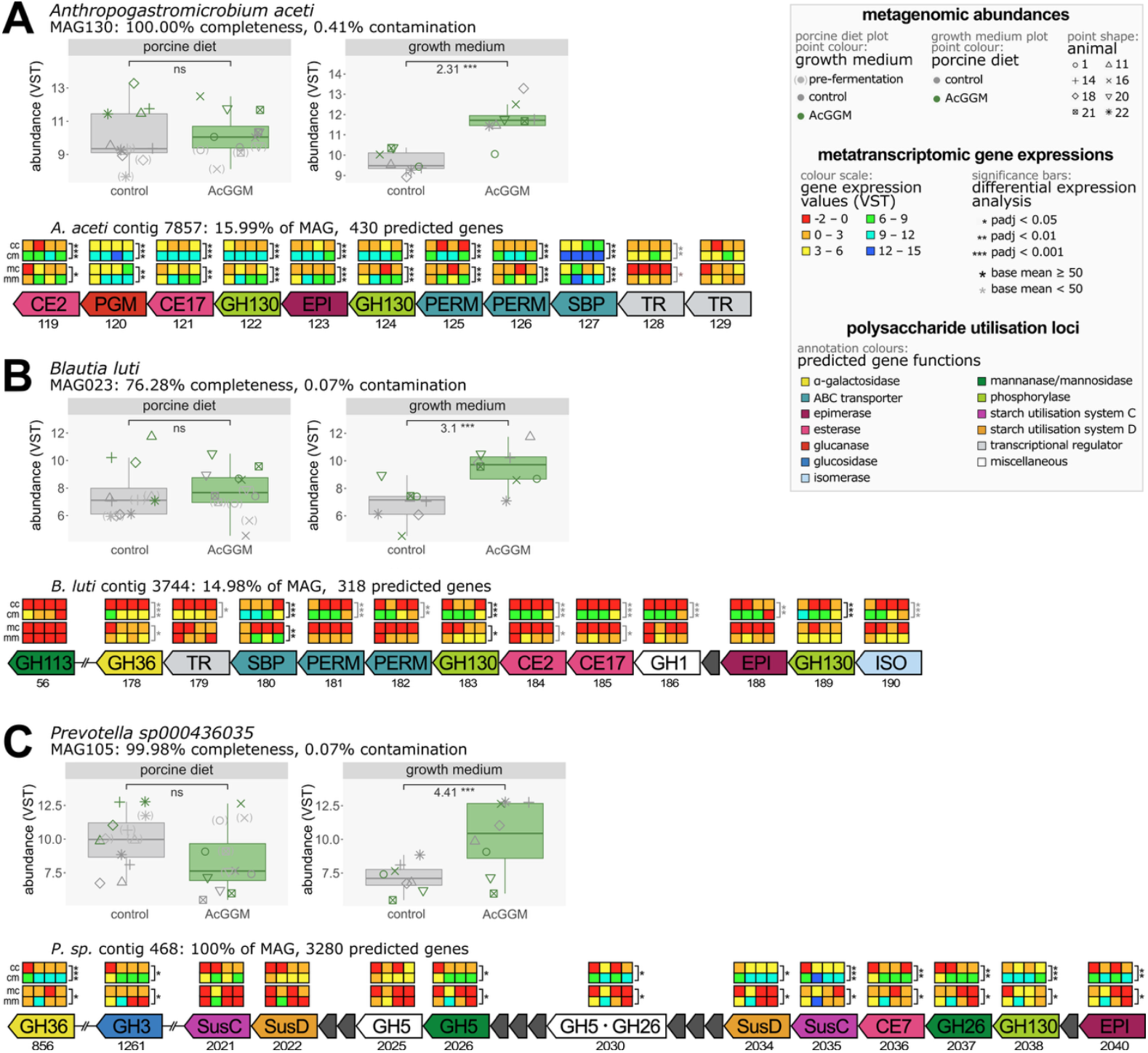
Metagenomic and metatranscriptomic data for three populations of interest, classified as **A)** Anthropogastromicrobium aceti; **B)** Blautia luti; and **C)** Prevotella sp000436035. **Upper panel**: boxplots showing variance-stabilised population abundances in samples belonging to the control or the AcGGM-supplemented group with respect to either porcine diet (left) or in vitro growth medium (right), with differential abundance test results displayed using significance bars atop each box pair (thresholds for significance: absolute LFC>1, base mean>50, and FDR p-value<0.05 (*), <0.01 (**), or <0.001 (***). None of the three populations displayed significant differences (ns) in samples derived from pigs in different diet groups, but all experienced an increased abundance when grown in the presence of AcGGM in vitro. **Lower panel:** putative mannan PULs annotated with predicted gene functions and variance-stabilised gene expression levels across the four total groups (cc: controls in porcine diet and growth medium; cm: control diet and AcGGM growth medium; mc: AcGGM-supplemented diet and control medium; and mm: AcGGM-supplemented diet and growth medium). Results of differential expression analyses within each subgroup are displayed as vertical bars using the same significance thresholds as described previously, in addition to grey annotations for significant differences when not thresholding the base mean.

*A. aceti* and *B. luti* both belong to the Bacillota phylum and have similar mannan PULs (**Fig. 4AB**) to those observed in other Bacillota members (12,14). Both lacked an extracellular endomannanase (GH26 or mannan-active GH5 subfamilies; 7, 8, 10, 17, 25, 36, 40, 41, 55, and 57) found in primary mannan degraders, hence they appeared only to be able to utilise mannooligosaccharides or behave as secondary degraders for polymeric mannan. Furthermore, both *A. aceti* and *B. luti* PULs had the ABC transporter proteins (SBP/PERM/PERM), two phosphorylases (GH130), two carbohydrate esterases (CE2 and CE17), an epimerase, and transcriptional regulators. In addition, *B. luti* had an upregulated α-galactosidase (GH36) and a mannosidase/mannanase (GH113) outside the PUL that was not upregulated. *A. aceti* did not have a GH36 in this PUL, although several GH36s were detected in other contigs of the MAG; however, no metatranscriptomic data were captured for these loci. The mannan PUL in *Prevotella sp000436035* (**Fig. 4C**) contained the carbohydrate uptake system SusC/D, GH26 endomannanases, GH5 mannanases/mannosidases, a phosphorylase (GH130), an esterase (CE7), and an epimerase. Additionally, an upregulated α-galactosidase (GH36) and glucosidase (GH3) were found outside the PUL. A characterised CE7 from a *Bacteroides* sp., which belongs to the same phylum as *Prevotella*, has shown activity on AcGGM (34). Similarly, another recent study on *Segatella copri* (previously known as *Prevotella copri*) revealed that a CE7 is actively involved in mannan degradation (35). These CE7s could not completely deacetylate mannan, indicating that *Prevotella* species, as well as *S. copri*, may only partially deacetylate mannans; which is perceived to be a disadvantage compared to Bacillota species containing a CE2-CE17 esterase pair.

Taken together, these observations indicate that *Prevotella* sp. – with its GH26 – may act as a primary degrader of AcGGM. *A. aceti* and *B. luti* may grow on the mannooligosaccharides existing in the AcGGM sample or cross-feed on shorter oligosaccharides produced by the extracellular endomannanase activity of other bacteria, similar to the cross-feeding previously seen between members of Bacteroidota and Bacillota (14). The cell surface-attached carbohydrate-binding protein in *Roseburia intestinalis* has been shown to bind oligosaccharides with a higher affinity than that reported for Bacteroidota species (12,36). If this strong binding affinity for oligosaccharides is common among Bacillota species, it may partly explain how Bacillota can function as secondary degraders by utilising oligosaccharides released by endomannanases from other species in a microbial consortium.

In addition to the three aforementioned putative AcGGM-degrading populations, there were 19 other MAGs whose abundance differed significantly after being inoculated with or without added AcGGM fibres in InVitSim (**Suppl. C**). Of these, 13 increased and 6 decreased in abundance (**Fig. SC3**). Several Lachnospirales appeared to perform better following *in vitro* supplementation of AcGGM; they included the Blautias *B. difficilis* (LFC>2) and *B. fusiformis* (LFC>1), and *Coprococcus spp*. (LFC>2). Population increases in AcGGM-grown communities also occurred for six different *Prevotella* strains, including *P. copri* (LFC>1) and *P. faecis* (LFC>2). On the other end of the spectrum, three Clostridia (two uncultured and one *Hominenteromicrobium*) and two Bacilli (*Enteromonas spp*. and *Bulleidia spp*.) (all LFC<-1) were less abundant when grown in AcGGM-supplemented medium. Among the genes that showed increased expression in taxa with higher abundance in AcGGM-supplemented fermentors, several CAZymes known to be involved in AcGGM degradation had higher expression levels: ABC transporters, CE families 2; 7; and 17, GH families 5; 26; 36; and 130, as well as cellobiose epimerases and phosphoglucomutases. (16, 23, 34, 36). The observed increase in abundance of these taxa when AcGGM was present *in vitro* alludes to their active fibre utilisation.

To supplement the metagenomics and -transcriptomics observations, we measured the concentration of key SCFAs in InVitSim following the *in vitro* fermentation in the presence or absence of AcGGM fibres. Acetate, propionate, and butyrate were all detected at higher concentrations in AcGGM-supplemented fermentors (**Fig. SD1**). Furthermore, the relative prevalence of these SCFAs across experimental groups correlated well with abundances of microbial populations with differential abundance – and thereby expression levels (**Suppl. A**) – following *in vitro* AcGGM supplementation (**Fig. 5**). All populations with higher abundance in AcGGM-supplemented *in vitro* growth medium showed positive correlations with observed SCFA levels, including the three putative AcGGM-degrading populations highlighted earlier in this section. While some of these microbes – like the *Prevotella sp*. – may directly contribute to increased SCFA concentrations through AcGGM degradation, others – such as *A. aceti* and *B. luti* – may act as secondary degraders and have their growth facilitated by the fibre degradation activity of other populations. Nevertheless, there is a prominent connection between these AcGGM-promoted microbes and the increased concentration of these SCFAs. Moreover, populations with limited growth in AcGGM-supplemented InVitSim fermentors negatively correlated with SCFA levels. Further research is needed to ascertain whether this trend is due to the accumulation of SCFAs, or the inability of these taxa to compete with those that can utilise AcGGM or its derivatives. Still, the consistency between the two omic layers was prominent and underlines the utility of considering multiple omic data layers for *in vitro* study approaches.

**Figure 5.**
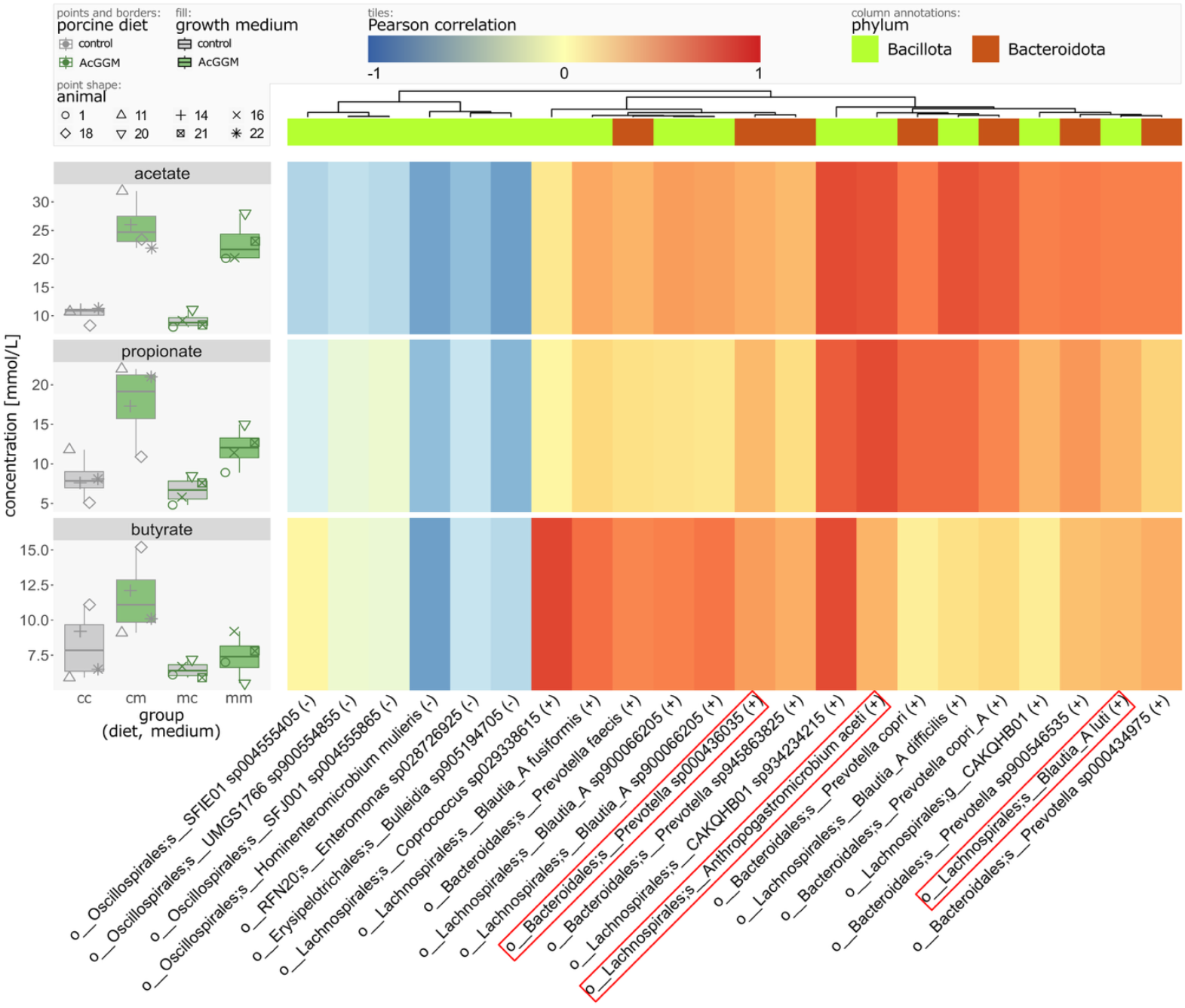
Concentrations of the short chain fatty acids acetate, propionate, and butyrate across sample groups (left), and Pearson correlation of these acid concentrations with population abundances (right). All taxa displayed were observed to be significantly differentially abundant between control and AcGGM-supplemented growth medium samples. Correlation values range from blue (−1) to red (1). Columns are clustered using Euclidean distance and Ward D2 criterion, and annotated with the phylum of each respective population. Taxon names below the heatmap inform on order and the most specific classification available for the population, and whether it was found in lower (-) or higher (+) abundance when AcGGM was provided in the growth medium. The three taxa of particular interest in the present study have been highlighted with red boxes. A full overview covering six measured short-chained fatty acids and all identified MAGs is included in **Suppl. D**.

To expand upon the observed effect of AcGGM on the microbial communities – both *in vivo* and *in vitro* – we conducted a principal component analysis (PCA) on each of the omic data layers: metagenomics reflecting the samples’ microbial compositions (**Fig. 6A**); and metatranscriptomics representing the transcriptionally realised activity of each community (**Fig. 6B**).

**Figure 6.**
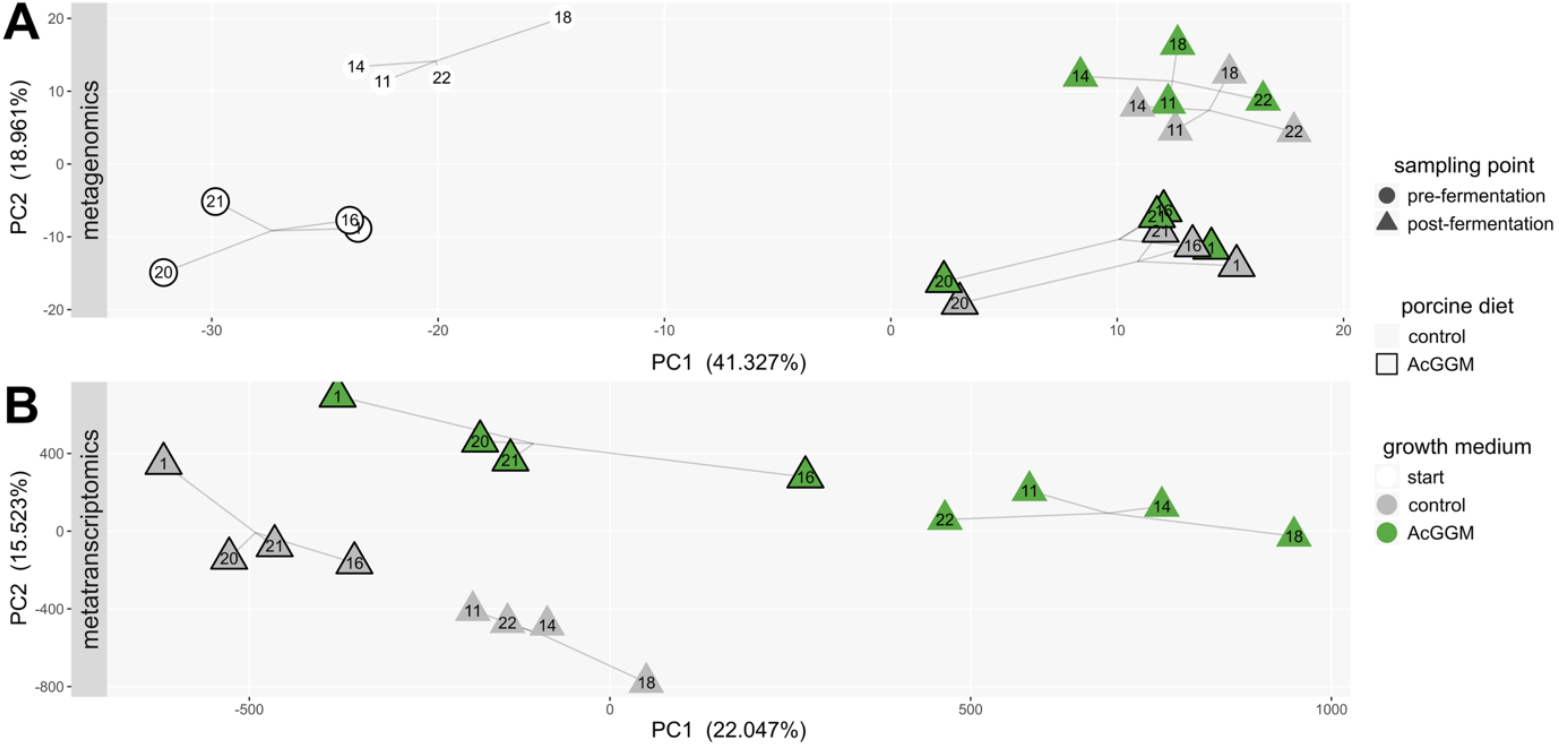
Principal component analysis of omic layers. Points represent samples with labels identifying donor animals. Line segments between sample subsets meet at the mean coordinate for each group. Axis labels report percentage of explained variance for each component. **A)** Variance-stabilised MAG abundances across samples. The first component captured a distinction between microbial communities present in samples before and after fermentation, while the second component saw a separation between samples derived from animals in different diet and growth medium groups. **B)** Variance-stabilised gene expression values. The components yielded distinct clusters of the four sample subgroups and indicated an effect of AcGGM-adaptation of the in vitro community due to the greater distance between the two groups of AcGGM-grown samples from different porcine diet groups (green), compared to control medium-grown groups (grey).

The first two components fit to either data layer yielded distinct clusters of samples that aligned with the experimental variables. Whether samples were collected pre- or post-fermentation appeared to be the most important factor in explaining microbial abundance patterns across samples (**Fig. 6A**). From inoculation and until the exponential growth phase, the microbial community compositions in the fermentors shifted similarly irrespective of porcine diet and growth medium. The impacts of porcine diet and *in vitro* growth medium with or without AcGGM were instead captured by the second component and showed that samples largely maintained their relative placement on the second axis irrespective of their growth medium, indicating that the *in vivo* AcGGM supplementation had a greater influence on the microbial composition than the *in vitro* fibre addition. While the *in vitro* AcGGM supplementation also caused shifts along both axes, the direction of which only matched that of the *in vivo* fibre administration along the first axis. Hence the impact of AcGGM on the microbial community composition appeared to only partly correspond between the *in vivo* and the *in vitro* model. Furthermore, the prior exposure of the microbiota to AcGGM fibres is alluded to as a factor that affected the microbial community composition. The functional omic layer (**Fig. 6B**) strengthened this impression by displaying a greater distance along the first axis between samples of different diet groups grown in AcGGM-supplemented medium *in vitro* compared to controls. Hence not only was the microbial community composition affected by the AcGGM supplementation, but their gene expression patterns also indicated that microbes altered their activity in response to the fibre. Moreover, this difference in transcriptomic activity was greater between AcGGM-adapted than non-adapted communities.

To corroborate our interpretations of the PCA, we identified populations and genes whose abundance and expression levels differed the most across sample groups through differential abundance and expression analyses, respectively. We investigated the effect of each key experimental variable in the present study by dividing the samples into groups according to sampling time, porcine diet, and growth medium. Detailed findings from the differential abundance analyses are included in **Suppl. A**; significant results from the metagenomic analyses have been visualised in **Suppl. C**; and all differential abundance and expression analysis results have been included in the GitHub repository accompanying the present work (jennymerkesvik/3domics_wp7_in-vitro-fermentation). Briefly, the results of these analyses further demonstrated that sampling time was the most impactful factor on the observed microbial communities (159 differentially abundant MAGs: 60 up and 99 down after fermentation), followed by porcine diet (78 total: 47 up and 31 down in AcGGM), and lastly growth medium (22 total: 16 up and 6 down in AcGGM). With respect to porcine diet groups, most observed changes (84% of instances) occurred within the Bacillota phylum. AcGGM fibre inclusion yielded a shift from Lachnospirales to other Clostridia. Taxa with increased abundance in samples from AcGGM-fed animals also displayed almost exclusively increased transcriptomic levels, indicating that their prevalence was due to increased metabolic activity; for instance a changed theoretical capacity for SCFA biosynthesis (**Fig. 2C**). While the upregulated genes encoded few AcGGM-degrading enzymes, several other CAZymes were upregulated in samples from AcGGM-fed animals. These observations suggest that promoted taxa may degrade other carbohydrates present in the porcine diet, such as xylan and starch, or act as secondary degraders of the AcGGM fibre.

To highlight the changes in microbial abundance and expression patterns that occurred irrespective of the growth medium used in InVitSim, the discussion thus far has not differentiated between communities extracted from animals given the control diet and those already introduced to AcGGM during the 4-week feeding trial. Since the preadaptation to AcGGM *in vivo* seems to impact the relative abundances of populations in the caecal microbiota (**Fig. 6**), extracted communities may have had different starting points for their continued development when inoculated in InVitSim. A primary AcGGM-degrader could, for instance, be outcompeted by populations utilising other carbon sources from the control diet and thus be at a disadvantage before AcGGM is introduced in a subsequent *in vitro* fermentation. Hence, we next investigated the sample subsets *within* the overall “porcine diet” contrast – namely the sample groups **cm** and **mm** (**Fig. SC4**) – to assess the effect of *in vivo* adaptation to AcGGM prior to fibre supplementation *in vitro*.

There were 76 MAGs with significant differential abundance between samples from animals of different diet groups when grown in the presence of AcGGM *in vitro*, 15 of which did not appear as such in the broader “porcine diet” comparison. These populations thus represent taxa whose *in vitro* growth in the presence of AcGGM is affected by whether they came from pigs fed AcGGM or not *in vivo*. Three Ruminococcaceae (*R. callidus* and *sp000433635* with LFC<-4, *Gemmiger qucibialis* with LFC<-1), two Lachnospiraceae (*Agathobacter spp*., *Fusicatenibacter saccharivorans*, LFC<-3), and a *Prevotella* (*P. sp945863825*, LFC<-3) displayed suppressed growth in AcGGM-supplemented media when extracted from animals fed the same fibre compared to the control diet. Conversely, three Eubacteria (uncultured, LFC>6), two other Clostridia (*Blautia difficilis, Gemmiger spp*., LFC>1), two Gammaproteobacteria (*E. coli, Mesosutterella spp*., LFC>4), and a Negativicutes (*Megasphaera elsdenii*, LFC>1) performed better in communities adapted to AcGGM prior to inoculation in InVitSim with the fibre supplement. The metatranscriptomic data showed significant differences when comparing samples from control- and AcGGM-fed animals grown in AcGGM for all but four of these populations. However, none of the genes with increased expression levels encoded prominent AcGGM-degrading enzymes. The observed advantage these populations had *in vitro* does not seem to be due to their ability to degrade the AcGGM fibre alone. Rather, the utilisation of other nutrients – including other carbohydrates and secondary metabolites from other taxa – could be the key to their prevalence. For instance, *B. difficilis* – which experienced improved growth when previously exposed to AcGGM – had increased expression of genes encoding non-AcGGM-degrading carbohydrate-active enzymes when AcGGM-adapted populations were inoculated in fibre-supplemented medium *in vitro*. The diminished *Prevotella* population, however, displayed downregulation of several genes needed for AcGGM degradation, including a region containing CE7, GHs 5 and 26, SusC, and SusD in close proximity (34,37). This observation points to this *Prevotella* population as a potential AcGGM degrader that persisted in control conditions *in vivo* by utilising other nutrient sources. Once transferred to InVitSim and given the AcGGM-supplemented medium, the non-AcGGM-adapted population activated its fibre degradation machinery to utilise the new carbon source to a greater extent than the AcGGM-adapted population. This may be due to it being outcompeted by another population with greater AcGGM degradation potential, which had become dominant in the *in vivo* AcGGM supplementation.

While the overall effect of AcGGM supplementation appears to be consistent across the present *in vivo* trial and the InVitSim system for several key taxa (e.g., members of *Prevotella* and Lachnospiraceae), there are species- and strain-level discrepancies between the microbial community developments after AcGGM is provided as a feed additive versus in a laboratory setting. Mannan metabolism has been suggested as one of the core pathways in the human gut microbiota; consequently, many bacteria have mannan degrading machineries (38). However, notably many taxa have mannan degrading machineries that remain dormant (13). Therefore, taxa with observed AcGGM-degradation potential following *in vitro* fibre administration may not display the same functionality *in vivo* and vice versa. This is likely attributed to the complexity of the animal-microbiota system and interactions that occur between the host and microbiome. In addition, the slightly altered community composition following fermentation in InVitSim could be due to difficulties in culturing specific taxa. Candidate targets for MDF should thus not be based solely on their performance in *in vitro* trials. Nevertheless, fermentation experiments facilitate the study of carbohydrate degradation and the potential host services the involved microbial taxa may provide. Future *in vivo* feeding trials involving AcGGM or other complex carbohydrates may then be assessed with this knowledge in mind, guiding the continued exploration of the effect of MDFs on the microbiome and its implications on host health.

## Materials and Methods

### Substrates

AcGGM from Norway spruce was used as a supplement in the feeding trial at 4% inclusion level and as a carbohydrate source in the medium during *in vitro* fermentations. The AcGGM was produced in-house from dried wood chips following the protocol in Michalak *et al*. (13).

### Animals and feeding trial

The four-week feeding trial included 24 female piglets from six sows. The animals were randomly distributed into groups of four in six pens, in which two animals were assigned the AcGGM supplemented diet and two the control diet. In order to mitigate pen variation, the pigs were individually fed, allowing pigs with both diets to be in the same pen. After the feeding trial, the animals were euthanised by captive bolt stunning followed by exsanguination.

### Sampling and preparation of caecal samples

Samples from all piglets were taken at the endpoint of the feeding trial. For each animal, the gastrointestinal tract was immediately taken out following euthanasia. The pH in the cecum was measured while the material from the lumen of the cecum tip was collected. The samples were directly transferred to 50 mL Falcon tubes containing an anaerobic cryoprotective solution (20% w/v), vortexed, and frozen at −80°C until further use. The cryoprotective solution – containing sodium dihydrogen phosphate (6.0 g/L), sodium hydrogen phosphate (7.1 g/L), cysteine-HCl (1.0 g/L), riboflavin (0.3 g/L), sucrose (50 g/L), and glycerol (15% [vol/wt]) – was prepared and pH-adjusted to 6.8 before being sterile filtered into tubes and kept protected from light overnight in an anaerobic cabinet (Whitley A85 Workstation; Don Whitley, UK) in an atmosphere of 85% N_2_, 10% H_2_, and 5% CO_2_ (39).

### *In vitro* fermentation, “InVitSim”

To test the fermentability of AcGGM *in vitro*, caecal digesta samples from eight of the euthanised pigs were chosen: four that had been on a 4% β-mannan inclusion diet; and four from the control group. The fermentors were produced in-house and consist of a double-jacketed stainless-steel vessel with a lid made in PEEK, containing connection ports for gas, acid and base, and a gas-sealed septum sampling port. The fermentors were pH-controlled, had headspace continuously flushed with N_2_ gas, and were temperature controlled via circulation bath (Thermo Fisher Scientific Inc.) with external water flushing.

The fermentors were autoclaved, filled with sterile basal medium and supplemented with either AcGGM (0.5% w/v) or ddH_2_O (no additional carbohydrate source as control), and placed in an anaerobic cabinet overnight before inoculation. The modified basal medium contained the following ingredients: peptone water (2 g/L), yeast extract (2 g/L), NaCl (0.1 g/L), KH_2_PO_4_ (0.04 g/L), K_2_HPO_4_ (0.04 g/L), MgSO_4_·7H_2_O (0.01 g/L), CaCl_2_·6H_2_O (0.01 g/L), NaHCO_3_ (2 g/L), hemin (0.01 g/L), vitamin K (10 µg/L), sodium glycocholate (0.5 g/L), sodium taurocholate (0.5 g/L), and L-cysteine hydrochloride (0.5 g/L) (20). The media was adjusted to pH 6.0, matching the average pH measured in the animals’ cecum at the trial end.

Samples were thawed inside the anaerobic cabinet and filtered through sterile 100 µm cell strainers (VWR International) before inoculation. The sample from each animal was divided into two fermentors, supplemented with either AcGGM or ddH_2_O. The digesta slurry samples were added to fermentors at 1% (v/v) final concentration and a total volume of 70 mL. After inoculation, the fermentors were taken out of the anaerobic cabinet, placed on magnetic stirrers, and attached to gas tubes (continues flushing of headspace with nitrogen (O_2_ < 5 ppm)), the pH control panels (peristaltic pumps with 1 M HCl and NaOH), and to a circulation bath holding 39°C (resembling the physiological temperature in pigs).

Samples for metagenomics and SCFA analysis were taken from the inoculation sample (timepoint 0 h) and during the exponential phase (timepoint 9 h). Samples for metatranscriptomics were taken during the exponential phase. 1 mL of samples for metagenomics and metatranscriptomics were mixed (1:1) with RNAlater (Sigma-Aldrich, USA).

### Metagenomics

Samples were sent to DNASense for sequencing and the generation of a MAG catalogue. Their described protocol was as follows: DNA was extracted using DNeasy PowerSoiul Pro Kit (QIAGEN, Germany), and concentration and purity were measured with Qubit dsDNA HS Assay and NanoDrop One (Thermo Fisher Scientific, USA). Barcoded SQK-NBD114.96 DNA libraries were prepared following manufacturer’s protocol (Oxford Nanopore Technologies, UK), loaded onto primed FLO-PRO114M (R10.4.1) flow cells, and sequenced with MinKNOW (R24.02.10) on a PromethION P24 device. Dorado Basecaller Server (ver. 7.3.9) (Oxford Nanopore Technologies, UK) was used for base calling and demultiplexing of signal data. Adapters were trimmed with Porechop (ver. 0.2.4) (40), and Nanoq (ver. 0.10.0) (41) was used to remove low-quality reads and produce sequencing data statistics. Draft *de novo* assemblies were produced with Flye (ver. 2.9.3-b1797) (42) using flags `-meta -extra-params min_read_cov_cutoff=10`. Medaka (ver. 1.11.3) (Oxford Nanopore Technologies, UK) was used to polish the drafts before inspection using Bandage (ver. 0.8.1) (43) and removal of contigs below 1000 bp with Seqkit (ver. 2.8.0) (44). Binary sequence alignment mapping files were created using Minimap2 (ver. 2.28-r1209) (45) with flags `-ax map-ont -I 8G -secondary=no` and Samtools (ver. 1.19.2) (46), and SemiBin2 (ver. 2.1.0) (47) with flags `-self-supervised -sequencing-type long_read -minfasta-kbs 500` was used to bin metagenome-assembled genomes. Dereplication was done using the dRep (ver. 3.5.0) (48) workflow with options `-comp 0 - con 10 -p -l 500000`, and CheckM2 (ver. 1.0.1) (49). MAG abundances were estimated using the CoverM (ver. 0.7.0) (50) genome workflow with flags `-p minimap2-ont -min-read-percent-identity 95 -min-read-aligned-percent 90`, before completeness and contamination levels were evaluated with CheckM2.

The MAGs provided by DNASense were classified against GTDB (ver. 2.4.0, R220) (51) and annotated on a high-performance computer cluster using DRAM (ver. 1.4) (52). The feature matrix was filtered for genome completeness (>50%), contamination (<5%), and presence in more than one sample, reducing the MAG catalogue from 1,064 to 301 populations. A phylogenetic tree of this catalogue was created using PhyloPhlAn3 (ver. 3.1.68) (53).

### Metatranscriptomics

Metatranscriptomic reads were generated by DNASense. A summary of the described protocol is given: RNA was extracted using the Monarch Total RNA kit (New England Biolabs, USA) following the manufacturer’s protocol, except bead beating at 6 m/s for 4*40 seconds. Product integrity and purity of RNA extracts were validated through gel electrophoresis using Tapestation 220 and RNA ScreenTapes (Agilent, USA). RNA concentrations were measured using Qubit RNA HS/BR Assay before DNA removal with TURBO DNAfree kit (Thermo Fisher Scientific, USA), and another round of quality assessment with RNA ScreenTapes and Qubit RNA HS/BR Assay. rRNA was depleted from the extracts with Ribo-Zero Plus rRNA Depletion Kit (Illumina, USA) and purification using the standard CleanPCR SPRI beads protocol (CleanNA, NL). Libraries were subsequently prepared using NEBNext Ultra II Directional RNA kit (New England Biolabs, USA). Concentrations were measured using Qubit HS DNA assay, and DNA size estimated using TapeStation with D1000 ScreenTape (Agilent, USA). Samples were pooled in equimolar concentrations and sequenced (2*151 bp, PE) on an AVITI platform (Element Biosciences, USA).

The cDNA reads from DNASense were quality-checked, pre-processed and filtered using FastQC (ver. 0.12.1) (54) and FastP (ver. 0.23.4) (55). Ribosomal RNA was removed from the dataset by mapping reads to the SILVA 16S/23S rRNA databases (`--ref silva-bac-16s-id90.fasta silva-bac-23s-id98.fasta silva-arc-16s-id95.fasta silva-arc-23s-id98.fasta`) using SortMeRNA (ver. 4.0) (56). Filtered reads were then quantified by pseudo-alignment to the MAG catalogue using Kallisto (ver. 0.46.1) (57). A total of 400,855 microbial genes were then mapped to the filtered catalogue of 301 MAGs.

### Short-chain fatty acids

were analysed as volatile fatty acids on GC-FID prepared according to a slight modification of method (58). Samples were prepared as follows: 1 mL sample was withdrawn from the fermentor and applied directly into a test tube, 4.25 mL of MeOH was added, and 0.8 mL C13:0 (0.5 mg/mL) was used as internal standard. 0.56 mL of 10 N KOH was added before tubes were capped and shaken vigorously for 1 minute on a vortex mixer, placed in a water bath at 55°C for 1.5 hours (tubes shaken vigorously 5 times during incubation), then cooled under running water. 0.464 mL of 24N H_2_SO_4_ was added, and tubes were placed in a water bath at 55°C for 1.5 hours (tubes shaken vigorously 5 times during the incubation period), then cooled in running water. For extraction, 2.4 mL heptane was added, tubes shaken vigorously for at least 1.5 minutes, centrifuged in a swing-out centrifuge (5 minutes at 3000 rpm, room temperature): approximately 1.5 mL of the heptane layer was transferred to 2 mL GC vials and cap. The samples were then analysed by GC set up as follows: Trace GC Ultra-system with FID (Thermo Fisher Scientific, USA), operated with Chromeleon 7.2 software (Thermo Fisher Scientific, USA): Rt-2560 GC Capillary Column, 100 m, 0.25 mm ID, 0.20 µm (Restek, Cat# 13198), Injector temp: 250 °C, Split injection (1:40 split ratio), Injection volume: 1 µL; Helium as carrier gas kept at 2,70 bar; with the following gradient 140°C (kept for 5 minutes) to 240°C with 4°C/min, FID detection set to 250 °C, 50 min runtime.

### Statistical analyses

All statistical analyses were conducted in R (ver. 4.2.1) (59). Both metagenomics and -transcriptomics datasets were compared across treatment groups using DESeq2 (ver. 1.38.3) (60) with default settings and significance thresholds set to base mean > 50, absolute log2 fold change > 1, and false discovery rate-adjusted p-value < 0.05. Both omics datasets were normalised by a variance-stabilising transformation before being used for additional analyses from the base R stats package, including principal component analyses and hierarchical clustering by Pearson correlation and Euclidean distance with Ward D2 criterion. The tool distillR (ver. 0.3.0) (61) was used to summarise DRAM-derived KEGG Orthology (62) annotations as qualitative functional capacities of the microbial community. Hill numbers were calculated using the hilldiv2 package (ver. 2.0.4) (63), spanning species richness (q=0) and Shannon-Weaver entropy (q=1) diversities; phylogenetic diversity by using the PhyloPhlAn-generated tree (q=1); and the aforementioned genome-inferred functional traits derived from distillR (q=1). Results were visualised using R packages ggplot (ver. 3.5.1) (64), ggtree (ver. 3.6.2) (65), ggtreeExtra (ver. 1.8.1) (66), and pheatmap (ver. 1.0.12) (67).

## Data availability

Metagenomic and metatranscriptomic sequencing data are available in European Nucleotide Archive with accession number PRJEB90225. Assembled MAGs are published on FigShare with DOI 10.6084/m9.figshare.28853753. Raw and pre-processed count matrices, metadata, and R code utilised for the analyses are available in the GitHub repository jennymerkesvik/3domics_wp7_in-vitro-fermentation.

## Acknowledgements

This study was funded by the European Union Horizon 2020 project “3D’omics” (101000309). The study also benefitted from the Norwegian Research Council infrastructure grants “Norwegian Biorefinery Laboratory” (270038), “FoodPilotPlant Norway” (296083), and “National network of Advanced Proteomics Infrastructure” (295910). LJL was supported by a faculty-funded PhD project (1205051076) and the Centre for Industrial Biotechnology (309558). We wish to thank the Livestock Production Research Centre at the Norwegian University of Life Sciences for housing and daily care of the animals, and for facilitating and contributing to sample collection. We acknowledge the Orion High Performance Computing Centre and the Threadripper station – set up by Carl M. Kobel for the Microbial Ecology and Multi-Omics group – both associated with the Norwegian University of Life Sciences, for computational resources that contributed to producing the research results reported within this paper.

## Author contribution statements

- Conceptualisation: LJL, PBP, BW
- Data curation: JM, RMS
- Investigation: LJL, ÖCOU
- Methodology: LJL, SLLR, TRH
- Formal analysis: JM
- Visualisation: JM
- Writing (original draft): JM, LJL, BW
- Writing (review and editing): JM, LJL, ÖCOU, RMS, SLLR, TRH, PBP, BW
- Supervision: SLLR, TRH, PBP, BW
- Funding acquisition: TRH, PBP, BW

